# G2VTCR: predicting antigen binding specificity by Weisfeiler-Lehman graph embedding of T cell receptor sequences

**DOI:** 10.1101/2025.04.29.651344

**Authors:** Zicheng Wang, Yufeng Shen

## Abstract

The binding of peptide-MHC complexes by T cell receptors (TCRs) is crucial for T cell antigen recognition in adaptive immunity. High-throughput multiplex assays have generated valuable data and insights about antigen specificity of TCRs. However, identifying which TCRs recognize which antigens remains a significant challenge due to the immense diversity of TCR. Here we describe G2VTCR (Graph2Vec-based Representation and Embedding of TCR and Targets for Enhanced Recognition Analysis), a computational method that uses atomic level graph embedding to predict TCR-antigen recognition. G2VTCR represents antigens and the third complementarity-determining region (CDR3) of TCR sequences using graphs, in which nodes encode atomic identities and edges encode chemical bonds between atoms, and then uses Weisfeiler-Lehman iterations to produce embeddings. The embeddings can be used for supervised classification tasks in TCR-antigen binding prediction and unsupervised clustering of TCRs. We evaluated G2VTCR using publicly available paired TCR-CDR3/antigen data generated by antigen-stimulation experiments. We show that G2VTCR has better performance in both classification and clustering than other embedding methods including pre-trained protein language models. We investigated the impact of Weisfeiler-Lehman iterations and the sample size of TCR CDR3 on classification performance. Our results highlight the utility of atomic level graphical embedding of immune repertoire sequences for antigen specificity prediction.

## Introduction

The recognition of peptide-MHC complexes by T cell receptors (TCRs) is pivotal for T cell activation and antigen specificity in adaptive immune responses. The knowledge of antigen specificity of T cells has a range of applications including cancer immunotherapies, vaccine development, and the understanding of autoimmune diseases (1-5). Experimentally determining TCR-antigen specificity relies on methods such as tetramer (1, 6) staining or in vitro stimulation assays (7), which are often labor-intensive, time-consuming, and difficult to scale for large datasets. Given the immense diversity of CDR3 sequences, estimated to range between 13^20^ to 14^20^ combinations, experimental sampling covers only a small fraction of possible TCRs. This challenge motivates the development of computational prediction models that evaluate TCR-peptide interactions across a broad range of sequences (8). Recent advancements in computational biology (9, 10)and graph-based machine learning (11, 12)have significantly enhanced the precision of such predictions.

The binding specificity of TCR-peptide is determined by TRA, TRB, peptide, and MHC (13). For CD8 T cell, the beta CDR3 sequence along with the V and J genes are often primary contributors (14). Previously published computational tools for analyzing TCR patterns and predicting peptide–TCR interactions can be categorized into three approaches. The first approach uses unsupervised clustering algorithms to identify antigen-specific binding patterns by grouping similar TCR CDR3 sequences. Tools such as TCRdist (15), GLIPH (16), and DeepTCR (17) follow this strategy. The second approach uses supervised machine learning methods, in which CDR3 and peptide sequences are represented using techniques like one-hot encoding, k-mers, BLOSUM or amino acid indices, such as TCRex (18), SETE (2), NetTCR (19). ERGO (20), pMTnet (21), TITAN (10). The third approach represents CDR3 and peptides using latent embeddings from language models trained on large-scale protein sequences, such as PanPep (9). However, most approaches face challenges in generalizing to unseen TCRs (22). This limitation may arise from their inability to effectively represent the structural and biophysical characteristics of TCR-antigen interactions.

Here we introduce G2VTCR (Graph-based Representation and Embedding of Antigen and TCR for Enhanced Recognition Analysis), a computational method that utilizes graph embedding techniques for high-dimensional modeling of TCR CDR3 and antigen sequences. G2VTCR constructs atomic-level graphs from antigen and TCR sequences, treating atoms as nodes and chemical bonds as edges, and extracts context-aware subgraph features using Weisfeiler-Lehman relabeling. This atomic-level graph embedding algorithm may capture biophysical properties of CDR3 and antigen peptides. We assessed the application of embeddings in predicting TCR-antigen binding as a supervised classification problem and clustering TCR sequences using unsupervised methods.

To ensure robust evaluation and reduce overfitting, we split the data into independent training and testing sets with non-overlapping TCR-antigen pairs. We optimized key hyperparameters, including the number of Weisfeiler-Lehman (WL) iterations (23, 24) and embedding dimensions, and evaluated performance using auROC. We demonstrate G2VTCR’s ability to accurately predict TCR-antigen interactions and its robustness across a variety of antigen pools (25).

## Results

### Model architecture

G2VTCR represents the sequences of both TCR CDR3 and antigens as graphs, in which nodes correspond to atoms (e.g., carbon, oxygen, nitrogen) and edges represent bonds between them. The method includes three main steps: (1) Seq2Mol, which converts sequence data into molecular structures; (2) Mol2Graph, which maps molecular structures into graph representations based on atomic and bond attributes; and (3) Graph2Vec (23), which generates graph embeddings using the Weisfeiler-Lehman graph kernel (26) (**Figure 1**). As part of the G2VTCR framework, the resulting graph embeddings are used for both supervised classification and unsupervised clustering. Specifically, a Random Forest (RF) classifier is employed to predict TCR-antigen binding, while DBSCAN is applied to identify clusters of TCR sequences with shared antigen specificity.

**Figure 1.**
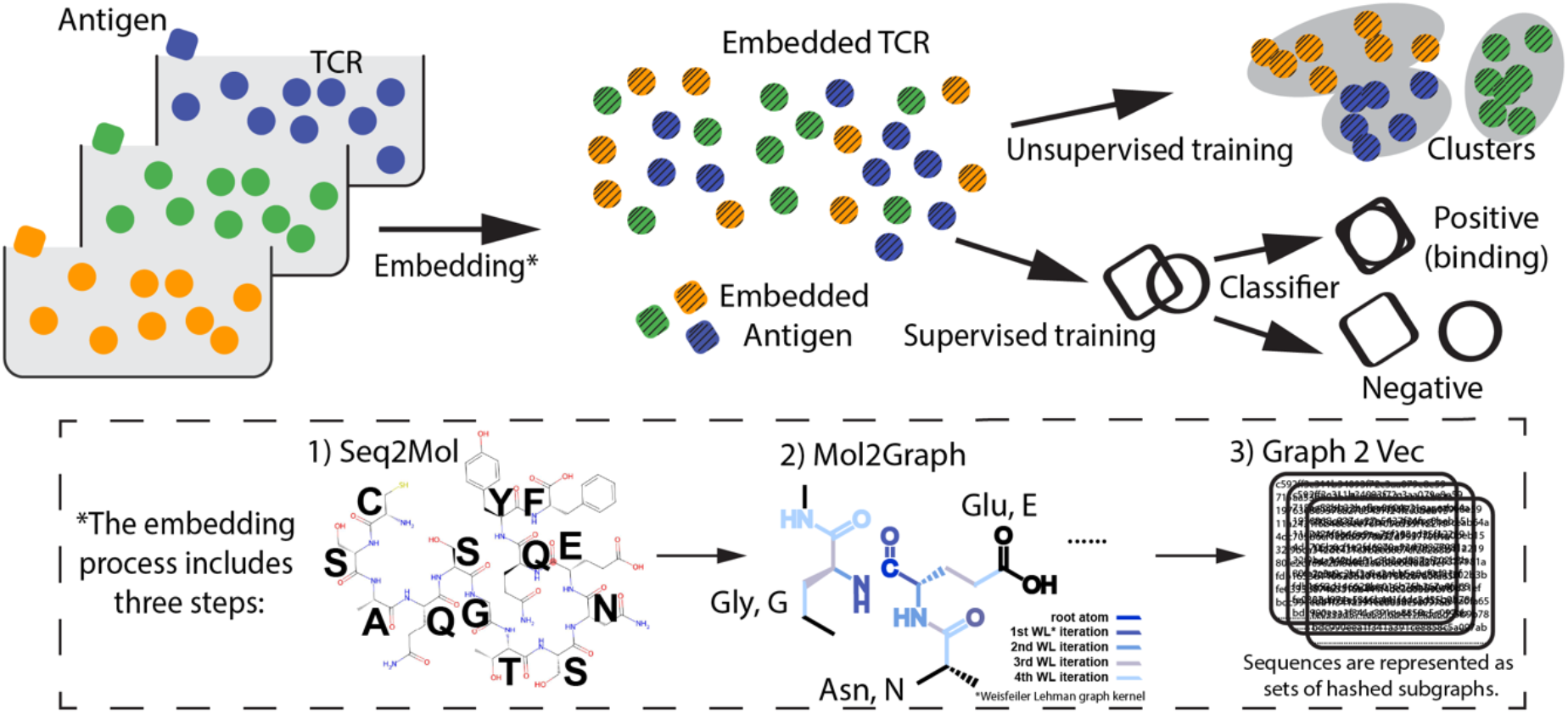
Schematic of the G2VTCR framework for predicting TCR-antigen binding using graph-based molecular representations. The method consists of three key steps: (1) Seq2Mol, where sequence data is converted into molecular structures; (2) Mol2Graph, where molecular structures are represented as graphs with nodes corresponding to atoms (e.g., oxygen, carbon, nitrogen) and edges representing bonds; and (3) Graph2Vec, where graph embeddings are generated using the Weisfeiler-Lehman (WL) graph kernel for iterative graph representation. The resulting TCR embeddings are applied to unsupervised clustering to identify groups of TCR sequences with similar antigen recognition profiles. Additionally, TCR and antigen embeddings are pooled together and used to train a Random Forest (RF) classifier, which distinguishes between positive (binding) and negative pairs.

We constructed training and testing dataset by pairing TCR CDR3 sequences with known antigenic peptides. Positive labeled pairs consisted of experimentally verified binding pairs, while negative pairs were generated through random sampling. The dataset was derived from two primary sources: (1) antigen stimulation experiments that identified TCR sequences from the COVID-19 research cohort (25). These data included expanded T-cell clonotypes, high-throughput sequencing of TCRβ CDR3 regions, and antigen specificity annotations; and (2) the VDJdb database (27), which provided a curated set of annotated TCR sequences with known antigen specificities.

As depicted in Figure 1, we pooled the TCR-antigen data and represented each sequence as a graph using the Seq2Mol and Mol2Graph steps. Using Graph2Vec, the method converted molecular interactions into numerical embeddings. The G2VTCR method then combined graph-based embeddings and machine learning classifiers to predict TCR-antigen interactions.

### Analysis of TCR Embeddings

To visualize TCR embeddings, we used t-distributed Stochastic Neighbor Embedding (t-SNE) to project the high-dimensional embeddings into two dimensions. We selected TCRs corresponding to 16 dominant epitope pools (Supplementary Table 1) to be shown in the figure. As shown in Figure 2, the t-SNE visualization revealed distinct groups corresponding to TCR sequences binding to different peptides, indicating that G2VTCR embeddings effectively capture antigen-specific characteristics.

**Table 1.** Performance Evaluation of Node and Edge Features in G2VTCR Framework. C1, C2, etc., represent different feature sets combining node and edge features for evaluating the G2VTCR framework.

|  |  |  |  |  |  |  |  |  |  |  |
| --- | --- | --- | --- | --- | --- | --- | --- | --- | --- | --- |
|  | C0 | C1 | C2 | C3 | C4 | C5 | C6 | C7 | C8 | C9 |
| Atomic Num |  | X | X | X | X | X | X | X | X | X |
| Is Aromatic |  | X | X | X | X | X | X | X | X | X |
| Formal Charge |  |  |  |  | X | X | X | X | X | X |
| Hybridization |  |  |  |  | X | X | X | X | X | X |
| Degree |  |  |  |  |  |  |  | X | X | X |
| Total Num Hs |  |  |  |  |  |  |  | X | X | X |
| Bond Type |  | X | X | X | X | X | X | X | X | X |
| Is Aromatic |  |  | X | X |  | X | X |  | X | X |
| Is Conjugated |  |  | X | X |  | X | X |  | X | X |
| Bond Stereo |  |  |  | X |  |  | X |  |  | X |
| Is In Ring |  |  |  | X |  |  | X |  |  | X |
| auROC | 0.951 | 0.966 | 0.965 | 0.963 | 0.964 | 0.963 | 0.961 | 0.954 | 0.955 | 0.955 |

**Figure 2.**
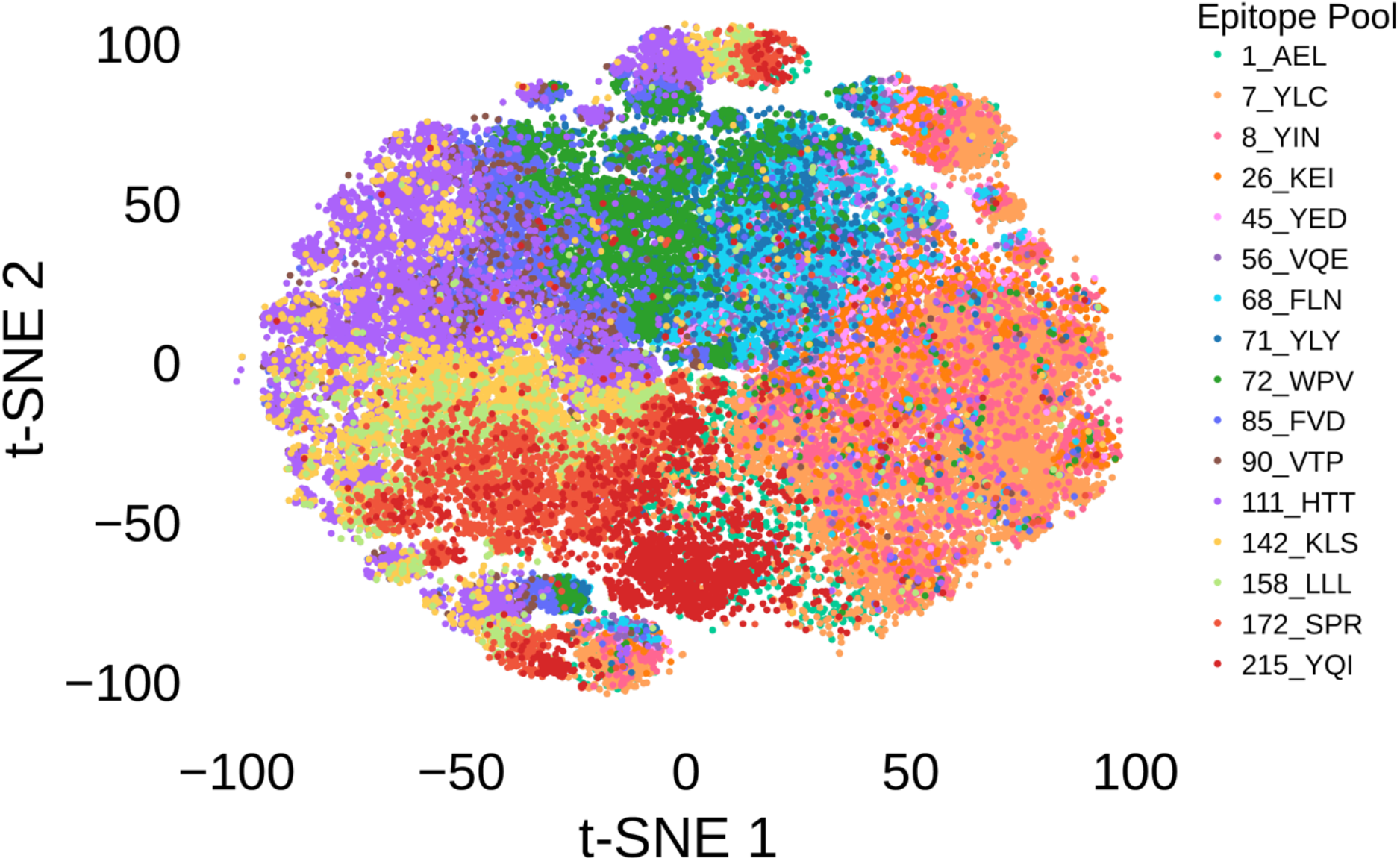
Embedding of TCR sequences. t-SNE visualization showing the distribution of TCR sequences across different epitope pools, highlighting their separation based on structural and antigen-specific features. Each color corresponds to a distinct epitope pool.

We selected antigens that are paired with at least 1,000 TCR CDR3 sequences for model training and testing. The TCR data were split into training and testing sets (80/20 split), ensuring that each TCR-antigen pair appeared exclusively in one set, with no within-epitope overlap. Instead of random split, we applied a sequence similarity-guided approach to construct the test set to minimize the similarity of TCRs between training and testing sets. Specifically, for each epitope group, we randomly selected seed TCRs and identified their nearest neighbors based on one-hot encoded sequence embeddings using FAISS with L2 distance. This KNN-like strategy ensured that similar sequences remained together, reducing the risk of data leakage and maintaining a clear separation between training and testing distributions (see Methods and Supplementary Figure 1). To generate negative samples, we used a shuffling strategy by pairing TCRs with peptides other than their known targets.

### Impact of Weisfeiler-Lehman Iterations, Embedding Dimensionality on classification

To identify optimal parameter settings, we performed parameter tuning using a 5-fold cross-validation scheme, where evaluation data was part of the training process and not used for testing. We examined the impact of key parameters in the Weisfeiler-Lehman procedure on the utility of the generated embeddings for classification. Specifically, we varied the number of WL iterations from 1 to 10 and the embedding dimensions across 16, 32, 64, 128, 256, and 512. A Random Forest classifier was trained using the embeddings produced by G2VTCR as input, with labeled data indicating TCR-antigen binding (positive interactions) or non-binding (negative interactions).

Graph2Vec embeddings were generated with dimensions ranging from 16 to 512, and model performance was evaluated using auROC scores across multiple Weisfeiler-Lehman (WL) iterations. A key factor in Graph2Vec’s success is the number of Weisfeiler-Lehman (WL) iterations, which determine the depth of neighborhood information incorporated into node labels. Fewer WL iterations capture local structures, while more iterations, performance gains diminish, and can lead to overfitting. Similarly, the dimensionality of embeddings influences how much information is encoded. High dimensions provide marginal improvements but may not justify the added complexity.

Figure 3 A and B show the area under the receiver operating characteristic curve (auROC) on 5-fold cross-validation as a function of Weisfeiler-Lehman (WL) iterations (1-10) across different embedding dimensions (16, 32, 64, 128, 256, and 512), highlighting how intermediate values offer the best balance between capturing structural complexity and avoiding overfitting. Our results indicate that an intermediate number of WL iterations (4-6) and a dimensionality of 512 provide the best performance for the given dataset. We further evaluated the robustness of G2VTCR embeddings when applied to RF classifiers with varying maximum depths. Notably, the performance in Supplementary Figure 3 corresponds to the scenario with unlimited maximum depth in the RF classifier, where an AUROC of 0.96 is achieved. When evaluated the maximum depth from 1 to 15, G2VTCR maintains consistently high performance after RF maximum depth equals to 7 (auROC > 0.9).

**Figure 3.**
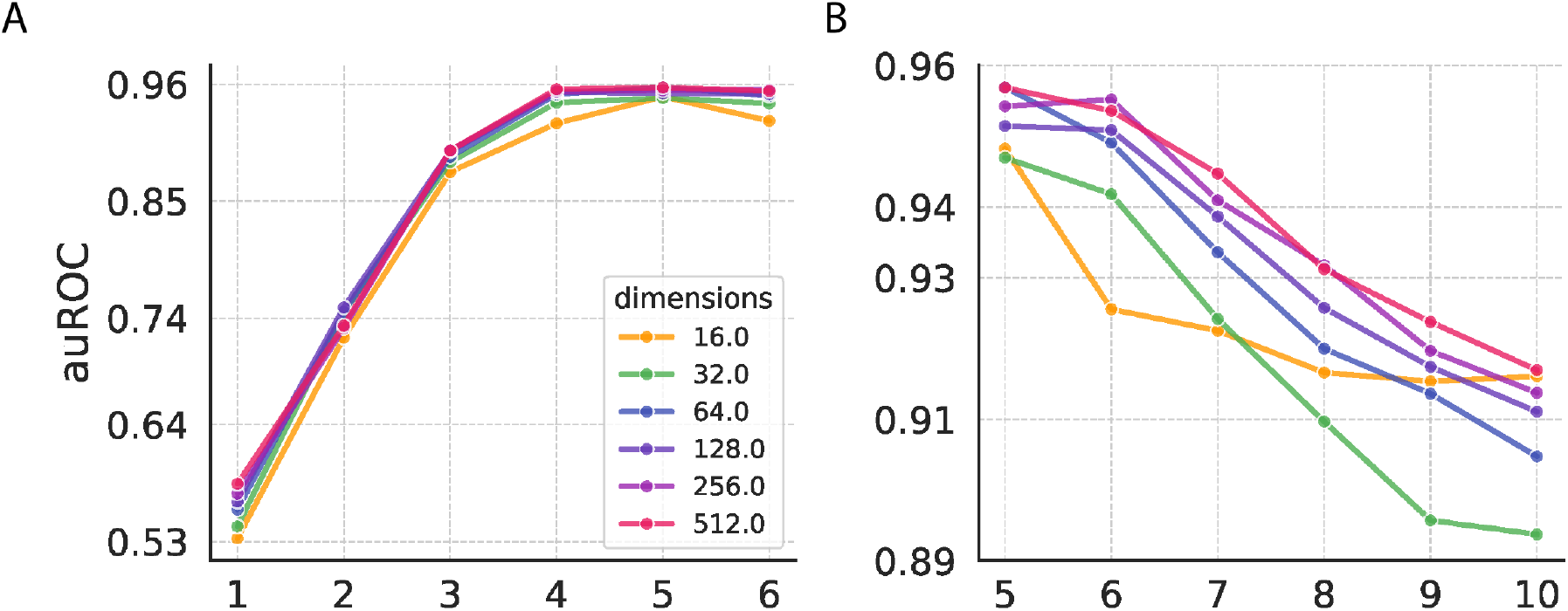
Impact of Weisfeiler-Lehman Iterations, Embedding Dimensionality. (A-B) Performance (auROC) on validation data as a function of Weisfeiler-Lehman (WL) iterations and embedding dimensionality, demonstrating optimal performance with 4-6 WL iterations and high dimensions (64+).

### Performance evaluation of classification and clustering based on G2VTCR embeddings

We next evaluated the predictive performance of G2VTCR embeddings in both supervised classification and unsupervised clustering tasks using test data. In the classification task, the model was trained to distinguish between positive TCR–antigen binding pairs and negative pairs generated by shuffling TCRs across non-cognate peptides. Each pair was represented by the concatenation of Graph2Vec embeddings of the TCR and epitope, learned from atomic-level graph representations. The Graph2Vec model was configured with five Weisfeiler-Lehman iterations and 256 embedding dimensions, selected based on prior cross-validation experiments. The classifier, a Random Forest, was trained on these combined embeddings using labeled training data and evaluated on a held-out test set constructed using a nearest-neighbor-based similarity splitting strategy to minimize overlap and leakage. Figure 4A shows comparative performance analyses revealed that the G2VTCR framework significantly outperformed other embedding methods in predicting peptide-TCR interactions. G2VTCR achieved an area under the receiver operating characteristic curve (auROC) of 0.96 for testing data. In contrast, the traditional One-hot encoding approach yielded auROCs of 0.71, while the more recent computational embedding method, ESM2 (650 M) (28), achieved lower auROCs of 0.61.

**Figure 4.**
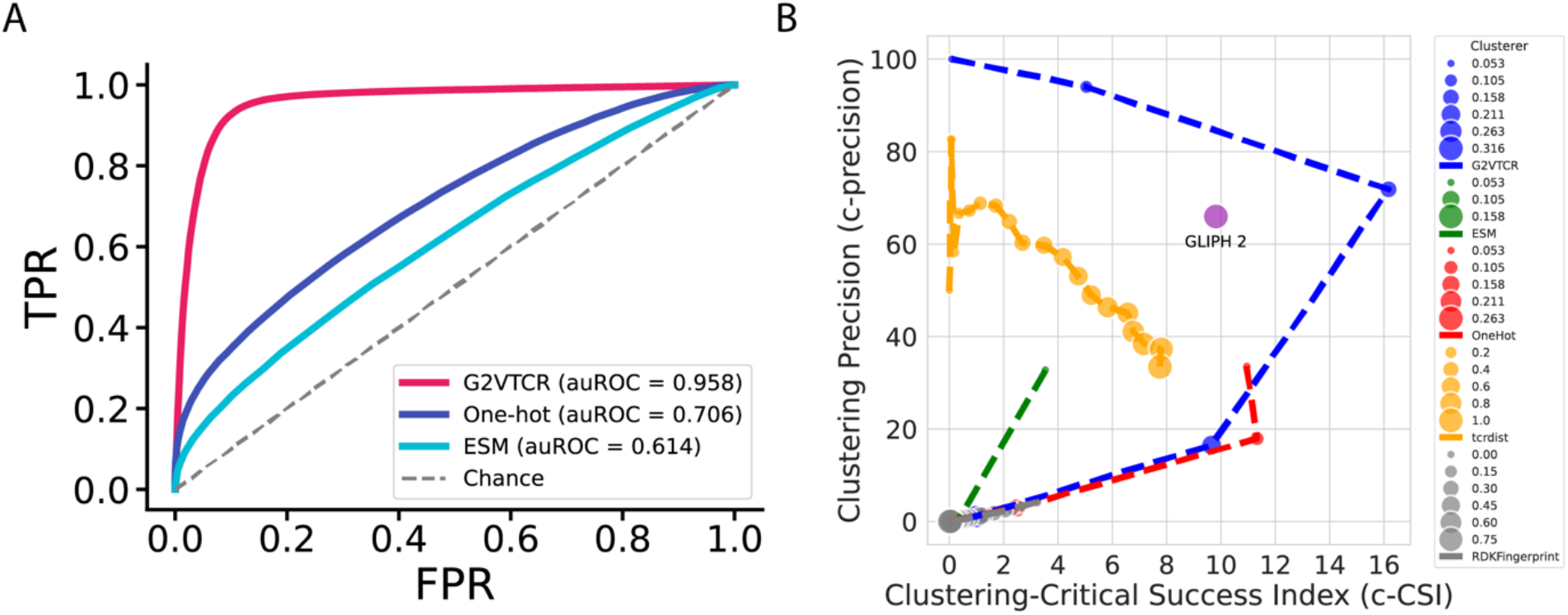
Receiver Operating Characteristic (ROC) curve of classification analysis and Clustering analysis of TCR sequence representations. (A) Shows performance under the shuffling strategy, with G2VTCR again achieving the highest auROC of 0.958, compared to One-hot (auROC = 0.706) and ESM2 (auROC = 0.614). (B) Clustering performance comparison across representations using Clustering Precision (c-precision) and Clustering-Critical Success Index (c-CSI) metrics across varying DBSCAN epsilon values. Representations include G2VTCR, ESM, OneHot encoding, RDKit fingerprint embeddings, and TCRdist. The G2VTCR embedding demonstrates superior performance, achieving the highest clustering precision and c-CSI, indicating its effectiveness in differentiating TCR clusters.

We evaluated the performance in clustering based on G2VTCR embedding. To construct the clustering dataset, we excluded TCR sequences associated with antigens represented by fewer than 1,000 unique sequences, thereby focusing the clustering analysis on antigens with sufficient representation. Using DBSCAN as the clustering algorithm, we varied the epsilon (*ε*) parameter to control the density threshold for cluster formation. Clustering performance was quantified using Clustering Precision (c-Precision) and the Clustering-Critical Success Index (c-CSI) (29). As shown in Figure 4B, the G2VTCR framework achieved higher clustering precision and c-CSI compared to alternative embeddings, including ESM2, OneHot encodings, RDKit fingerprints (30), TCRdist embeddings (15) and GLIPH 2 (16).

To evaluate whether the similarity between test and training data affects model performance, we computed the average minimal TCR distances by first calculating the pairwise distances between test and training CDR3 sequences for each epitope using the TCR-dist method (15), and then averaging the minimum distance for each test sequence with training sequence from the same epitope pool (Supplementary Figure 4). We found there was no significant correlation between the average minimal TCR distances and the auROC, as indicated by a Pearson correlation coefficient of 0.12 and a p-value of 0.49, suggesting a weak and non-significant relationship. Futhermore, there is no significant correlation between the number of TCRs and the auROC (correlation coefficient=0.014, p-value=0.94). Epitope-specific analyses nonetheless highlight the robustness of the G2VTCR framework, with several antigens demonstrating exceptional predictive performance. Notably, antigen pools 37, 141, 90, and 215 achieved auROC scores of 0.989, 0.986, 0.984, and 0.981, respectively (see Supplementary Table 1 for sequences and indices).

### Impact of data depth on performance

To further investigate the impact of data depth on model performance, we conducted down-sampling analysis on individual epitopes from the COVID-19 research cohort dataset (31), analyzing the relationship between the number of TCR sequences (Figure 5A) and the data ratio (Figure 5B) with the corresponding auROC scores. To simulate varying data availability, we applied down-sampling prior to the embedding step, after splitting the data into training and testing sets. In each experiment, both the training and testing sets were down-sampled at the same ratio to ensure consistency. These analyses reveal how data availability influences predictive accuracy. As shown in Figure 5B, the strong positive correlation (*r* = 0.65, *p* = 4.06 × 10^*−*43^ r=0.65, p=4.06×10 ^*−*43^) indicates that increasing the number of TCRs generally enhances predictive performance, as measured by auROC scores. However, for epitopes with highly diverse repertoires, even datasets with over 1,000 TCRs may not fully capture all binding scenarios. Figure 5B shows the relationship between the data ratio (proportion of available TCR data) and model performance. The results reveal a critical threshold at a 20% sampling data, where predictive accuracy sharply increases, indicating sufficient data coverage for reliable predictions (Supplementary Figure 2). Beyond this point, performance gains become incremental, with diminishing returns observed after 30%.

**Figure 5.**
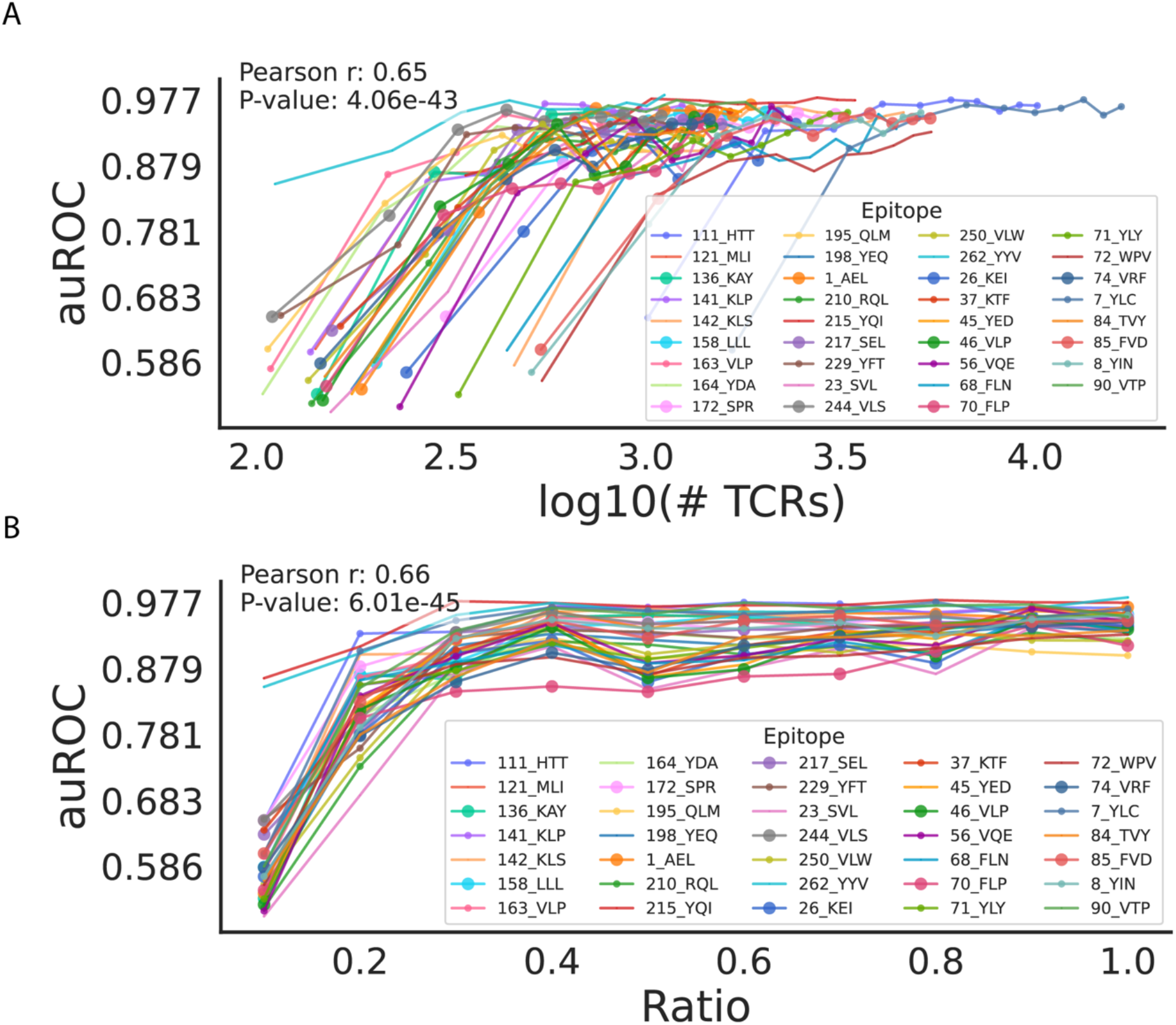
Data Depth on TCR-Epitope Interaction Prediction Performance. (A) Correlation between the number of TCRs (log10-transformed) and auROC scores for epitope-specific prediction, highlighting improved performance with increasing TCR counts. (B) Relationship between data ratio and auROC scores, showing an elbow point at 20% and a plateau beyond 30%, with epitope-specific variability in performance trends.

We observed a similar trend on clustering performance with the depth of the dataset used during the embedding step. Specifically, down-sampling the input data resulted in a noticeable decrease in clustering performance by down-sampling to 20% or below of the original data, as reflected by reductions in both clustering precision and c-CSI metrics (Supplementary Figure 2). This degradation affected G2VTCR and other embedding methods alike, leading to less distinct separation of TCRs into clusters, particularly within epitope-specific groups. These findings highlight the importance of adequate data coverage when using unsupervised embeddings to capture the diversity and specificity of TCR repertoires.

### Impact of Node Features and Subgraph Structure on Model Performance

We systematically evaluated the influence of various node and edge features on the performance of the G2VTCR framework in predicting antigen-TCR interactions. Node features such as atomic number, aromaticity, formal charge, hybridization, connectivity degree, and the total number of hydrogens were assessed alongside bond-level features including bond type, aromaticity, conjugation, stereochemistry, and ring membership. Performance was quantified using the area under the receiver operating characteristic curve (auROC), with a baseline of 0.951 achieved using a non-attributed model (wl_iterations=5, dimensions=128, embeddings based solely on graph structure and connectivity, without incorporating node or edge attributes).

Our findings suggest that the performance of the G2VTCR framework is robust across various feature configurations, with only minimal differences observed when specific features are added or excluded (Table 1).

### Identification of High-Frequency Motifs in TCR Sequence Clusters

We analyzed TCR sequence clusters based on their atomic-level subgraph patterns using the G2VTCR framework and TF-IDF (Term Frequency-Inverse Document Frequency) vectorizer (32)to the graph features of the sequences. This method allowed us to identify the most significant words or motifs common in the sequences within each cluster. We specifically focused on atomic features that appear frequently within the cluster sequence graphs but not across all clusters, ensuring these features are distinctive for each cluster. Here, we present the analysis of three enriched TCR clusters, highlighting key features and prominent patterns in the TCR sequences as examples.

In the high-accuracy prediction epitope pool 215_YQI, we identified several key patterns in the TCR sequences occurring at significant frequencies. The most prominent patterns included “SIGQG” (223 sequences), “SIGTG” (207 sequences), and “SIGLG” (174 sequences), all sharing the common “SIG” prefix, which represents a recurring motif within this cluster. Variations such as “SLGQG” (137 sequences), “SIGVG” (102 sequences), and “SMGQG” (90 sequences) highlighted flexibility around the core “SIG” and “SLG” motifs. Additional motifs, including “STGTG” (73 sequences), and “SSGTG” (35 sequences), further demonstrated the diversity of subgraph patterns within this cluster. We compared G2VTCR against GLIPH using real motif outputs from 215_YQI epitope pool, GLIPH identified frequent short patterns such as “IG” (1053 sequences), “SI” (1031), and “SIG” (982), which correspond to common substrings but lack positional specificity or structural insight.

To better illustrate how these motifs define cluster specificity, we extended our analysis to examine high-frequency motifs with high TF-IDF scores across additional epitope pools, including 7_YLC, 111_HTT. Figure 6 provides a logo plot visualization of these key motifs, capturing the probabilistic occurrence of specific patterns within the sequence clusters. This analysis highlights not only the distinctiveness of certain motifs within individual epitope pools but also their variation across clusters. These high-TF-IDF motifs, such as “%GAI%”, “SARGG” in epitope pool 7_YLC and “%GPWD”, RD/GP” in epitope pool 111_HTT, which GLIPH returned longer but sparse motifs like “RGLAG%SYE” and “S%GGNE” (each seen in ∼15 sequences, ambiguous or gapped positions indicated by “%”), which include ambiguous or gapped positions. Similarly, GLIPH identified “SPRD” and “SLRD” in epitope pool 111_HTT as short, frequent motifs.

**Figure 6.**
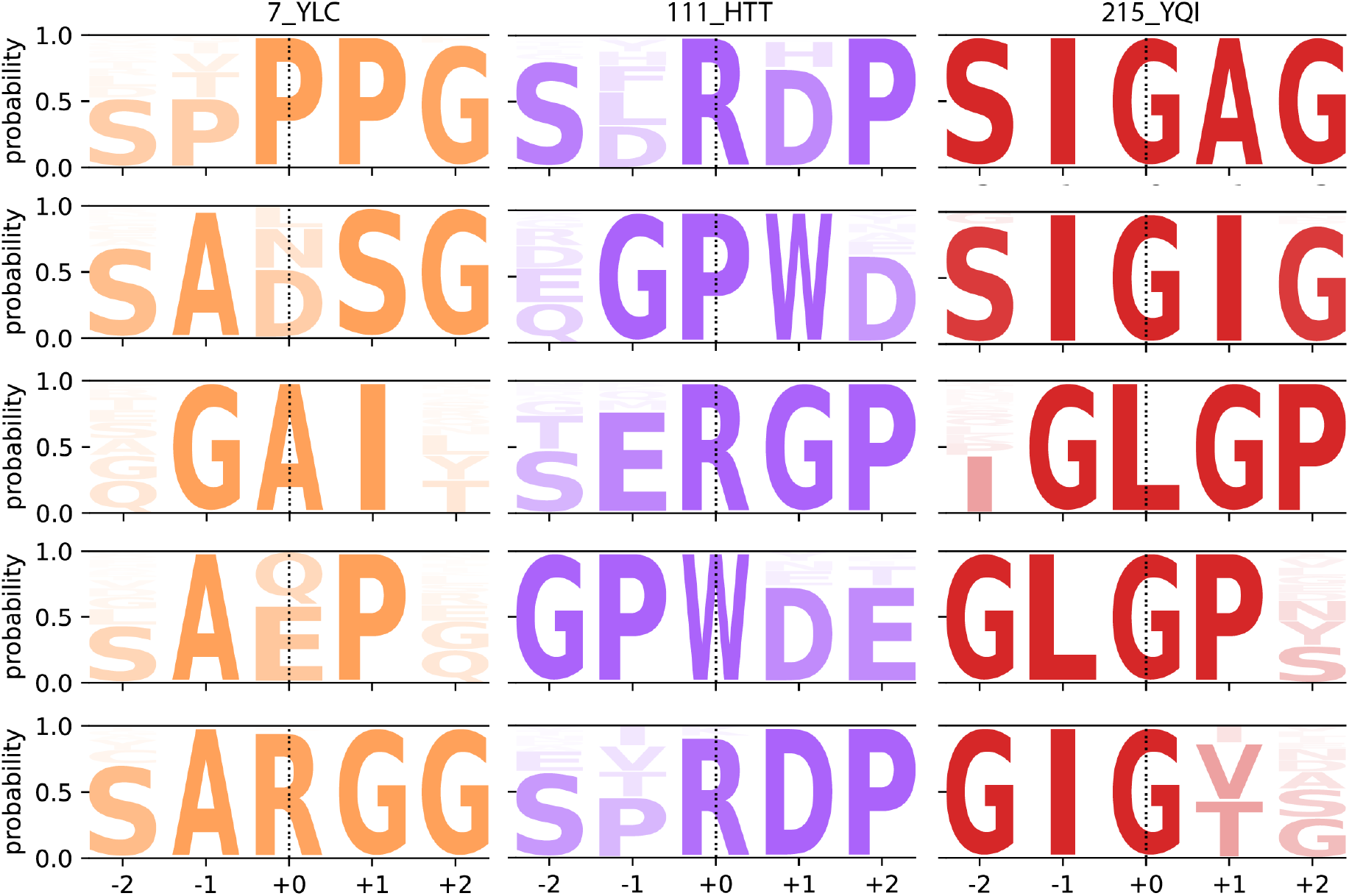
Sequence logo plots of high-frequency, high-TF-IDF motifs from TCR sequences across epitope pools 7_YLC, 111_HTT, and 215_YQI. Each motif is shown as a five-residue window centered at the most informative position (position 0). The number of motifs per epitope varies depending on the number of specific subgraph-based patterns that met the selection criteria: high TF-IDF scores, presence in at least 5 TCRs, no overlap with motifs from other epitopes, and occurrence in at least 20 TCR sequences in the dataset.

## Discussion

G2VTCR represents a new approach for modeling of TCR-antigen interactions. Leveraging atomic-level graph-based embeddings, G2VTCR achieves superior performance in both supervised classification and unsupervised clustering tasks compared to previous amino acid-level and language model-based embedding methods. This improvement may be attributed to its ability to more efficiently represent discontinuous binding motifs at the atomic scale. Unlike amino acid-level embeddings, which are limited by the coarse granularity of a 20-character corpus, G2VTCR’s atom-level embeddings extend the available information by capturing subgraph-level structural and chemical properties. This approach allows the model to preserve molecular details that are critical for accurately modeling TCR-antigen interactions. While sequence-based models such as OneHot and ESM2 are highly effective for longer protein sequences, they do not account for the local biophysical and chemical properties that are vital for short sequences like CDR3 regions.

Our analysis highlights the importance of optimal WL iterations and embedding dimensions in enhancing model performance. Intermediate WL iterations (4-6) and a dimensionality of 128-256 were found to provide the best balance, capturing detailed neighborhood information without introducing excessive noise.

Unlike k-mer-based methods that rely on fixed-length substrings for pattern detection (33), G2V embeddings operate at the graph level, enabling the identification of structurally equivalent patterns despite sequence variation. While k-mer methods, such as SPAN-TCR, can detect frequent motifs like “GX” or “XG” and calculate entropy reduction, they are limited by exact sequence matching and fixed k-mer sizes. In contrast, G2V embeddings use graph-level abstractions to capture inter-atomic interactions and represent TCR structural diversity comprehensively. For instance, the recurring “XGLGP” motif and its variants highlight not only shared atomic subgraph contexts but also the central residues critical for binding specificity, which k-mer-based approaches cannot discern due to their reliance on fixed-length substrings.

A limitation of G2VTCR is that the method’s predictive accuracy depends on comprehensive sampling of TCR sequences specific to each antigen. The performance of G2VTCR declines when applied to antigens for which only limited TCR sequence information is available. Additionally, our current evaluation did not fully explore the model’s capacity to generalize to novel antigens with sparse TCR representation. Future studies should address this potential limitation to confirm the robustness of G2VTCR across broader antigenic contexts.

In this study we have focused on TCR β-chain without considering α-chain or HLA alleles. This approach would not work well for class II–restricted T cell responses, where both α- and β-chain participate antigen recognition. Future studies could address this issue by incorporating paired α-β TCR sequences and multiple HLA alleles in larger and more diverse datasets.

## Acknowledgements

We thank Dr. Donna Farber, Dr. Peter Sims, and Jean-Baptiste Reynier for insightful discussions, and Jake Hagen for assistance with computing infrastructure. This work was supported in part by the National Institutes of Health grant U19AI128949.

## Methods

### Code Availability

Our implementation of G2VTCR is publicly available at https://github.com/princello/G2VTCR.

### Data Collection and Process

We used a dataset derived from antigen stimulation experiments to identify T-cell receptor (TCR) sequences from the MIRA assay of COVID-19 subjects (31). We included data on T-cell clonotype distributions, high-throughput sequencing of TCRβ CDR3 regions, and antigen specificity annotations obtained through computational analyses. The dataset originally contains 152,718 TCRs. After filtering out epitopes with fewer than 1,000 associated TCRs, 97,216 TCRs remained for the analysis. Before filtering, there were 545 epitopes from 269 antigen groups. After removing epitopes with fewer than 1,000 associated TCRs, 120 epitopes from 35 antigen groups remained for analysis. Additionally, we incorporated TCR sequences from the VDJdb database (27), which provided annotated TCRs with known antigen specificities.

#### Training and testing splitting

The dataset consisted of TCR CDR3 sequences paired with antigenic epitopes, divided into training and testing sets using an 80/20 split. To construct the testing set, we employed a nearest-neighbor sampling approach based on sequence similarity. Specifically, TCR sequences were one-hot encoded and embedded into a fixed-length vector space, and L2 distance was used to identify clusters of similar sequences using the FAISS library. Seed sequences were randomly selected, and their nearest neighbors were grouped to form the test set. This procedure ensured that sequences associated with the same antigenic epitope were entirely allocated to either the training or testing set, preventing within-epitope leakage and maximizing independence between sets. Additionally, we introduced two strategies for generating negative TCR–epitope associations: random shuffling of known TCRs across non-cognate epitopes and controlled swapping of TCRs between different antigen groups, both of which were included in the model training and evaluation.

#### Shuffling Known Positive Pairs

This approach involves creating random TCR-epitope combinations by pairing each TCR with epitopes different from their known matches. The assumption here is that a TCR specific to one epitope is unlikely to be specific to a different, unrelated epitope. Due to the MIRA assay can identify what might be considered “negative pairs” as part of its methodology by accomplishing the process of sorting T cells into antigen-specific and non-antigen-specific populations after exposure to antigen pools. the shuffed antigen-TCR Pairs could be considered as negative data.

#### Graph Construction

Antigen and TCR CDR3 sequences were input as strings of amino acid codes and converted into molecular structures using RDKit (30), a cheminformatics toolkit. In this representation, each atom within the amino acids, such as carbon, nitrogen, and oxygen, was treated as a node. The edges between these nodes represent chemical bonds, detailing the molecular structure of the peptide.

Following the conversion, the molecular structures were transformed into detailed graph representations. Nodes in these graphs were enriched with attribute data reflecting the atomic properties such as atomic weight, electronegativity, and other relevant chemical characteristics. Edges, representing the chemical bonds between atoms, included attributes such as bond type (e.g., single, double, peptide bond) and bond order, which are crucial for understanding the molecular architecture of the peptides.

The molecular graphs generated in RDKit were then exported to NetworkX (34)to facilitate the application of advanced network analysis techniques. This integration allowed for a comprehensive analysis that goes beyond simple structural depiction, enabling the study of detailed interactions within the peptide sequences. NetworkX provided tools to apply various graph-theoretical algorithms that analyze the structural and functional characteristics of the peptides at an atomic level.

Graph embeddings were performed to transform this detailed atomic graph data into a numerical form suitable for computational models. Techniques such as node-level, edge-level, and graph-level embeddings were employed to derive vector representations that capture the intrinsic chemical and structural properties of the peptides for predictive modeling tasks.

#### Graph Embedding with Graph2Vec

In the graph embedding process used by Graph2Vec, each graph *G* = (*V, E*), where *V* is the set of nodes and *E* is the set of edges, is represented as a vector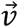. The algorithm treats each graph as a document and the rooted subgraphs around each node as words. For each graph *G*, the rooted subgraph *h* (*u*) for a node *u* is constructed by aggregating information from its *k*-hop neighborhood, where *k* represents the Weisfeiler-Lehman (WL) subtree height. The subgraph information is labeled and transformed into a bag-of-words representation BoW (*G*) using the WL subtree kernel.

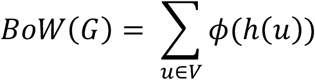

Here, *ϕ*(*h*(*u*)) represents the encoding of the rooted subgraph around node *u*.

The Graph2Vec algorithm then applies a document embedding technique to learn a fixed-size vector representation that captures the structural and label information of these subgraphs. This embedding vector 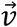 for each graph is obtained through an unsupervised learning approach that maximizes the likelihood of preserving neighborhood information of the graphs within the embedding space. This is achieved by optimizing the following objective function:

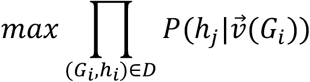

where 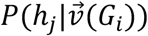 is the probability of the subgraph *h*_*i*_ given the graph embedding 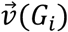, and *D* is the dataset of graphs. This probability is modeled using a softmax function:

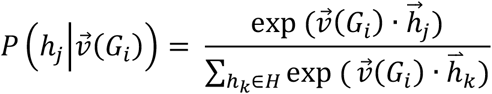

where 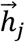 is the embedding of the subgraph *h*_*j*_, and *H* is the set of all subgraphs in the dataset. Through this unsupervised learning approach, Graph2Vec generates a vector 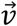 for each graph that captures both structural and label information of the subgraphs, preserving neighborhood information in the embedding space.

In addition to Graph2Vec, the embedding methods utilized in this study include OneHot Encoding, Evolutionary Scale Modeling (ESM), and RDKit Fingerprint Embedding. OneHot Encoding encodes amino acids as independent binary features. ESM generates sequence embeddings informed by evolutionary-scale protein databases. RDKit Fingerprint Embedding represents molecular graphs as binary vectors based on substructure presence. These methods were employed to compare their performance in capturing relevant features for downstream analyses.

#### TCR Distance Calculation

For the pairwise distance calculation between T-cell receptor (TCR) sequences, we used the TCR-dist method (15), which computes similarity based on the complementarity-determining region 3 (CDR3). This method relies on a vector-based distance metric that accounts for both the sequence and structural properties of the CDR3 regions.

Key parameters for the TCR-dist calculation included the use of a gap penalty of 4 to penalize insertions and deletions, and the trimming of 3 residues from the N-terminus and 2 residues from the C-terminus of each CDR3 sequence. The computation was optimized through the use of parallel processing across 4 CPU cores, and redundant sequences were filtered to improve efficiency. To further accelerate the calculations, we utilized precomputed distance matrices with appropriate weighting applied to the sequence distances.

This configuration ensured accurate and efficient pairwise distance estimation between TCR sequences, forming the basis for subsequent analyses such as clustering and visualization.

#### Performance metrics

Numerous benchmark methods have been proposed to assess the performance of TCR-epitope prediction algorithms, including those founded on sequence distance. True positive rate (TPR), false positive rate (FPR), and accuracy (ACC) are derived from the counts of true positives (TP), true negatives (TN), false positives (FP), and false negatives (FN). The area under the receiver operating characteristic curve (AUROC) is a metric to evaluate our classification model’s performance, gauging the model’s capacity to differentiate between positive and negative classes.

#### TF-IDF Score

TF-IDF (Term Frequency-Inverse Document Frequency) vectorization is a technique used to convert text data into numerical representations. It helps identify the importance of words (in this paper, hashed atomic labels) in a document relative to a collection of documents (graph). Here’s how it works and why it’s useful in identifying key features within clusters:

The term frequency (TF) measures how frequently a word appears in a document. It is calculated as:

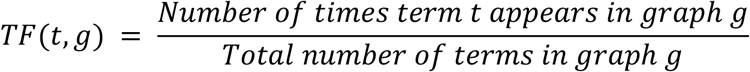

The invere document frequency (IDF) measures how important a word is in the corpus. It is calculated as:

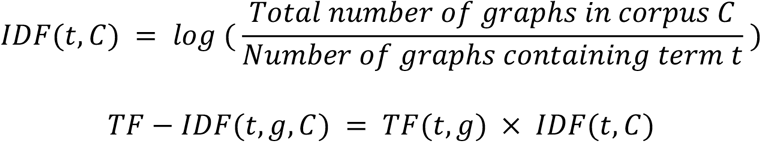

This score highlights hashed atomic features that are important (i.e., frequent in a particular graph but rare across the corpus). Additionally, the code implementation considered a minimum document frequency to filter out less significant features. Only features appearing in a minimum number of documents were included, ensuring that the identified features were not only significant within individual graphs but also commonly represented across multiple graphs in the cluster.

#### Clustering Analysis

To evaluate the clustering performance of various TCR sequence representations, we applied the DBSCAN (Density-Based Spatial Clustering of Applications with Noise) algorithm (29). DBSCAN was selected for its capability to detect clusters of arbitrary shapes while effectively handling noise. The algorithm relies on two key parameters: *ε*, which defines the maximum distance between points within the same neighborhood, and *m*, the minimum number of points required to form a cluster. For a given point *p*: The *ε*-neighborhood is defined as:

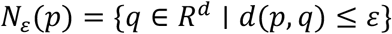

Where |*N*_*ε*_(*P*)| *≥ m* for *p* to be a core point.

A point *q* is density-reachable from a core point *p* if there exists a chain of core points *p*_1_, *p*_2_, *…, p*_*n*_ such that *p*_1_ = *p, p*_3_ = *q*, and *p*_*i+*1_ *∈ N*_*ε*_(*p*_*i*_)

DBSCAN was applied with a range of *ε* values (0.01 – 20) using the Manhattan distance metric.

Dimensionality reduction was applied to ensure computational feasibility and optimize clustering performance. Principal Component Analysis (PCA) was used to reduce fingerprint embeddings, Graph2Vec, ESM2 (650M), and OneHot encoding to 24 dimensions. Multidimensional Scaling (MDS) was applied to TCR distance matrices computed using TCRdist to project the data into a lower-dimensional space. The performance of the clustering algorithm was assessed using two key metrics: Clustering Precision (c-precision), this measures the purity of clusters concerning epitope labels. For a cluster *C*_*i*_

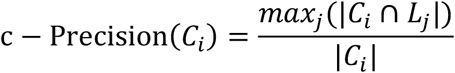

Where *L*_*j*_ represents the set of points with lable *j*, and |*C*_*j*_| is the size of cluster *C*_*i*_. Clustering-Critical Success Index (c-CSI): This metric accounts for both cluster purity and the proportion of well-clustered points:

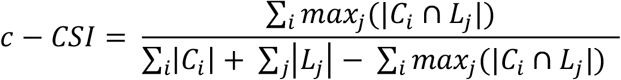

while c-CSI accounts for both cluster purity and the proportion of well-clustered points, providing a comprehensive evaluation of clustering quality. The clustering analysis was performed across the three primary TCR representations—graph-based embeddings, ESM2 embeddings, and OneHot encodings—as well as the RDKit-based fingerprint embeddings and TCRdist. The resulting metrics were plotted to compare the clustering effectiveness of each representation under varying levels of noise and clustering density. Since GLIPH does not involve varying *ε*-like parameters, the output of the clustering analysis is a single pair of c-Precision and c-CSI values for each processed dataset or condition. This is different from DBSCAN, where multiple metrics can be calculated across a range of *ε* values.

