## Supplemental Figures for "G2VTCR: predicting antigen binding specificity by Weisfeiler-Lehman graph embedding of T cell receptor sequences"

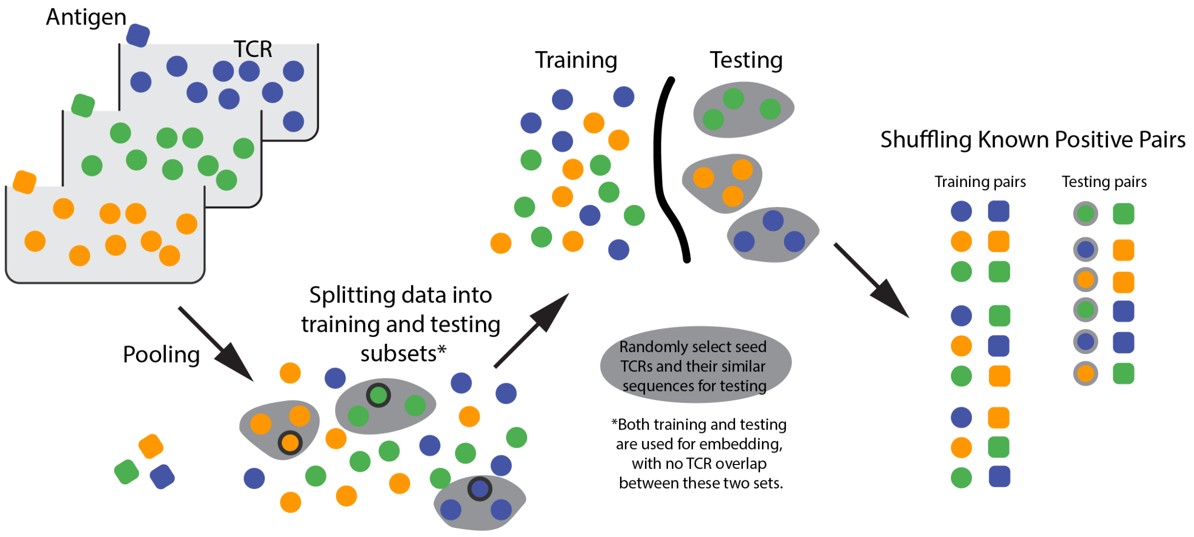


**Supplementary Figure 1.** Overview of Training and Testing Data Splitting Approach. The splitting process begins with TCR CDR3 sequences paired with antigenic epitopes. Data are divided into training and testing subsets, ensuring minimal overlap between the two. Testing subsets are generated by selecting sequences and their nearest neighbors based on sequence similarity, while training subsets consist of the remaining sequences.

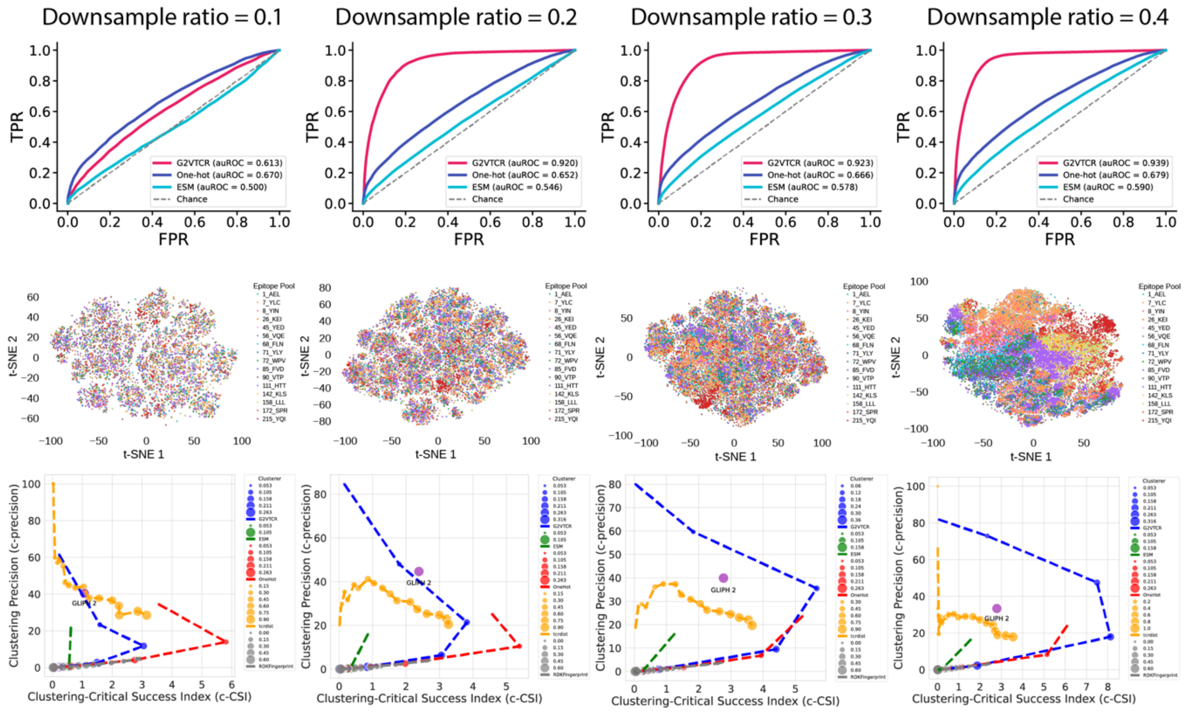


**Supplementary Figure 2**. Impact of Downsampling on TCR-Epitope Interaction Prediction and Clustering Analyses.
Each row corresponds to a different downsampling ratio (0.1 to 0.4), illustrating the effect of reduced data density on model performance and clustering analysis. Down sampling impacts both the predictive performance (first two columns) and the clustering metrics (last two columns).

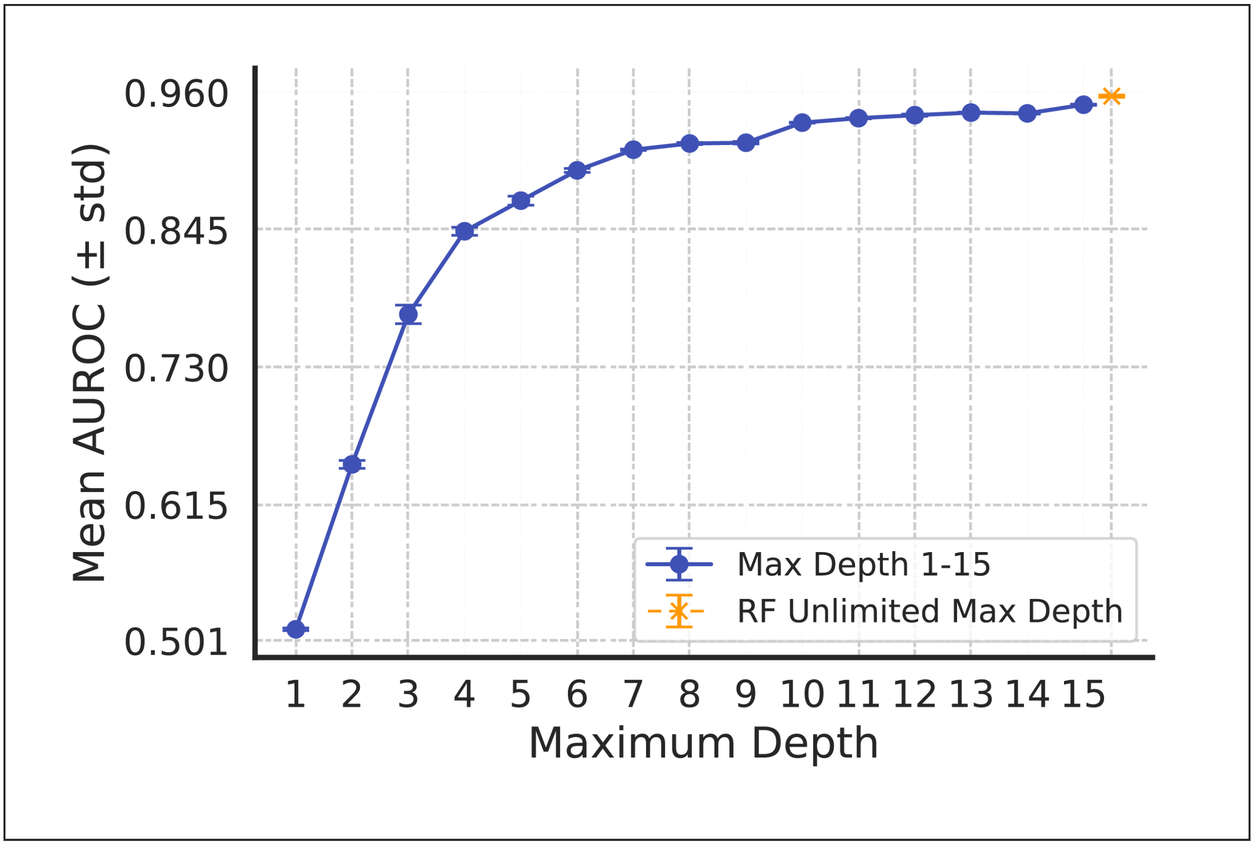


**Supplementary Figure 3.** Random Forest (RF) performance analysis for peptide-TCR interaction prediction. Mean AUROC (± standard deviation) of G2VTCR embeddings with RF classifiers across maximum tree depths (1–15). The performance stabilizes at deeper depths, with a maximum AUROC of 0.958 achieved with unlimited tree depth.

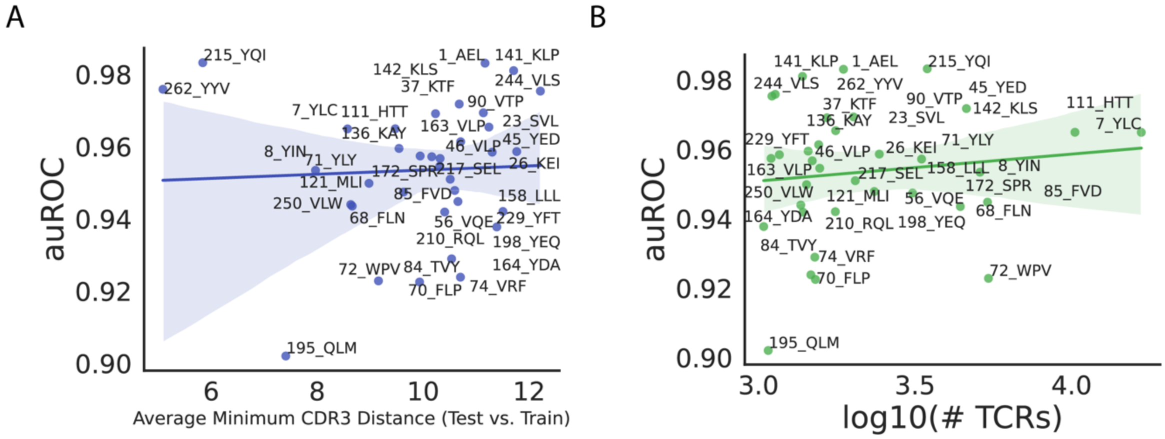


**Supplementary Figure 4.** The correlation between (A) the average minimal TCR distances of TCRs, (B) the number of TCR and auROC for epitope-specific prediction.

**Supplementary Table 1.** IDs, Amino Acid Sequences, ORF Coverage, and Genomic Coordinates of TCR-Antigen Associations.
